# A novel fully-human potency-matched dual cytokine-antibody fusion protein targets carbonic anhydrase IX in renal cell carcinomas

**DOI:** 10.1101/744029

**Authors:** Roberto De Luca, Baptiste Gouyou, Tiziano Ongaro, Alessandra Villa, Barbara Ziffels, Alessandro Sannino, Gianluca Buttinoni, Simone Galeazzi, Mirko Mazzacuva, Dario Neri

**Affiliations:** Philochem AG, Otelfingen, Switzerland; Department of Chemistry and Applied Biosciences, Swiss Federal Institute of Technology (ETH Zürich), Zurich, Switzerland; Philogen SpA, Monteriggioni, Italy

**Keywords:** Immunotherapy, antibody-cytokine fusion proteins, IL2, TNF, EDA domain of fibronectin, CAIX

## Abstract

Certain cytokines synergize in activating anti-cancer immunity at the site of disease and it may be desirable to generate biopharmaceutical agents, capable of simultaneous delivery of cytokine pairs to the tumor. In this article, we have described the cloning, expression and characterization of IL2-XE114-TNF^mut^, a dual-cytokine biopharmaceutical featuring the sequential fusion of interleukin-2 (IL2) with the XE114 antibody in scFv format and a tumor necrosis factor mutant (TNF^mut^). The fusion protein recognized the cognate antigen (carbonic anhydrase IX, a marker of hypoxia and of renal cell carcinoma) with high affinity and specificity. IL2-XE114-TNF^mut^ formed a stable non-covalent homotrimeric structure, displayed cytokine activity in *in vitro* tests and preferentially localized to solid tumors *in vivo*. The product exhibited a partial growth inhibition of murine CT26 tumors transfected for carbonic anhydrase IX. When administered to *Cynomolgus* monkey as intravenous injection, IL2-XE114-TNF^mut^ showed the expected plasma concentration of ~1500 ng/ml at early time points, indicating the absence of any *in vivo* trapping events, and a half-life of ~2 hours. IL2-XE114-TNF^mut^ may thus be considered as a promising biopharmaceutical for the treatment of metastatic clear-cell renal cell carcinoma, since these tumors are known to be sensitive to IL2 and to TNF.

**Contribution to the field:** There is a growing interest in the antibody-based targeted delivery of pro-inflammatory cytokines for tumor therapy, which may be complementary or alternative to immune checkpoint inhibitors for immunotherapeutic applications. In this article, we have described a novel fusion protein, featuring antibody moieties specific to carbonic anhydrase IX, as well as interleukin-2 and tumor necrosis factor as pro-inflammatory cytokines, which combines the ability to recognize a tumor-associated antigen on the surface of renal cell carcinomas with the simultaneous display of two potent therapeutic payloads. The newly developed product (termed IL2-XE114-TNF^mut^) exhibited favorable biochemical characteristics, the ability to preferentially localize at the tumor site, a cancer growth inhibition activity and a suitable pharmacokinetic profile in *Cynomolgus* monkey. IL2-XE114-TNF^mut^ may therefore represent an attractive candidate for the immunotherapy of renal cell carcinoma.

## Introduction

Antibody-cytokine fusions (also called “immunocytokines”) represent an emerging class of engineered cytokine products, that may display a superior anti-cancer activity as a consequence of a preferential accumulation at the tumor site, helping spare normal organs (1). An increased density of lymphocyte within the tumor mass typically correlates with a better prognosis, both in mice and in cancer patients (2–5). The targeted delivery of cytokines to the tumor environment may increase the intratumoral density of T-cells and NK cell (2,4,6). In this context, IL2 and interleukin-12 (IL12) have proven to be particularly attractive payloads for antibody-based delivery applications (7–9), as these agents can potently activate T-cells and NK cells.

Antibody-cytokine fusions with tumor-homing properties are likely to display their therapeutic action and to increase the therapeutic index of the corresponding cytokine payload as a result of a specific activation and proliferation of tumor-resident CD8+ T cells and of NK cells, which recognize malignant structures (2,3). For some pro-inflammatory cytokine payload (e.g., IL12), it has been shown that an antibody-based targeted delivery to the tumor may increase therapeutic activity by at least 20-fold, compared to the recombinant cytokine counterpart (10).

Antibody-cytokine fusion proteins in clinical trials for the treatment of cancer include various antibody-IL2 fusions (e.g., hu14.18-IL2 (11), huKS-IL2 (12), L19-IL2 (13), F16-IL2 (14), CEA-IL2v (15), NHS-IL2 (16), DI-Leu16-IL2 (17)), as well as fusions with TNF (e.g., L19-TNF (18)) and with IL12 (BC1-IL12 (19) and NHS-IL12 (20)). More recently, scientists at Glycart-Roche have described a novel antibody fusion with human 4-1BBL, which has started clinical trials (21). The most advanced product may be represented by a combination of L19-IL2 with L19-TNF, which is currently being investigated in Phase III clinical trials (EudraCT number: 2015-002549-72 and NCT03567889) for the treatment of fully-resectable Stage IIIB,C melanoma (22).

IL2 and TNF are representative examples of cytokines which work well when used in combination. Our group already described in 2010 that the simultaneous administration of L19-IL2 and L19-TNF in an immunocompetent mouse model of neuroblastoma was more active than the products given as single agents. Indeed, the combined use of the two products cured the majority of mice (23). A synergistic benefit for L19-IL2 and L19-TNF was also observed for intralesional administration procedures in mouse models of cancer (24). This work paved the way for the execution of a Phase II clinical trial in melanoma patients (22), which has then triggered Phase III study programs in Europe and in the United States. Tumor necrosis factor and IL2 are synergistic and complementary also from a pharmacodynamic viewpoint. TNF damages the tumor endothelium leading to a rapid hemorrhagic necrosis of the neoplastic mass (4,6,25), while IL2 mainly acts by activating NK cells and CD8+ lymphocytes (6,24,26).

The synergistic action of IL2 and TNF has stimulated research activities, aimed at incorporating both payloads into the same therapeutic product. We have previously described the production and anti-cancer activity of a dual-cytokine fusion protein (termed IL2-F8-TNF^mut^), which exhibited a selective accumulation at the tumor site following intravenous administration and a potent anti-cancer activity, particularly against murine soft-tissue sarcomas (6). The therapeutic action of IL2-F8-TNF^mut^ could be potentiated by combination with immune checkpoint inhibitors (27).

Here, were report the cloning, production and characterization (*in vitro* and *in vivo*) of a novel fusion protein (termed IL2-XE114-TNF^mut^), capable of recognizing carbonic anhydrase IX (CAIX) and of simultaneously displaying IL2 and a de-potentiated TNF mutant (TNF^mut^). CAIX is a membrane protein, which is overexpressed in hypoxia conditions and in various cancer types, including renal cell carcinomas (RCC), urothelial, colorectal, stomach, pancreas and other cancers (28,29). This antigen is virtually undetectable in most normal adult tissues, exception made for certain gastrointestinal structures (29). CAIX has been targeted *in vivo* using both antibody- and small molecule-based products, showing interesting results in imaging studies (30–32).

The product was active *in vitro* and *in vivo* and may represent a candidate for the immunotherapy of renal cell carcinoma.

## Materials and Methods

### Tumor cell lines

The human renal cell carcinoma cell line SKRC52 was kindly provided by Professor E. Oosterwijk (Radbound University Nijmegen Medical Centre, Nijmegen, the Netherlands). Transfected CT26-CAIX cells were prepared as previously reported (30). CHO cells, CTLL2 cells and L-M fibroblasts were obtained from the ATCC. Cell lines were received between 2017 and 2019, expanded, and stored as cryopreserved aliquots in liquid nitrogen. Cells were grown according the supplier’s protocol and kept in culture for no longer than 14 passages. Authentication of the cell lines also including check of postfreeze viability, growth properties, and morphology, test for mycoplasma contamination, isoenzyme assay, and sterility test were performed by the cell bank before shipment.

### Mice and tumor models

Six to eight-week-old female BALB/c nude mice were obtained from Janvier Labs. Tumor cells were implanted subcutaneously in the flank using 1×10^7^ cells (SKRC52), 3×10^6^ cells (CT26-CAIX). Experiments were performed under a project license (license number 04/2018) granted by the Veterinäramt des Kantons Zürich, Switzerland, in compliance with the Swiss Animal Protection Act (TSchG) and the Swiss Animal Protection Ordinance (TSchV).

### Cloning, expression and protein purification

The fusion protein IL2-XE114-TNF^mut^ contains the antibody XE114 (31) fused to a mutated version of human TNFα (arginine to alanine mutation in the amino acid position 108 of the human *TNF* gene, corresponding to the position 32 in the soluble form) at the C-terminus by a 15-amino acid linker and to human IL2 at the N-terminus by a 12-amino acid linker (6). The gene encoding for the XE114 antibody and the gene encoding human TNF and human IL2 were PCR amplified, PCR assembled, and cloned into the mammalian expression vector pcDNA3.1(+) (Invitrogen) by a NheI/NotI restriction site as described previously (6).

The fusion proteins used in this study were expressed using transient gene expression in CHO cells as described previously (33,34) and purified from the cell culture medium to homogeneity by Protein A (Sino Biological) chromatography.

### *In vitro* characterization

Purified proteins were analyzed by size-exclusion chromatography on a Superdex 200 increase 10/300 GL column on an ÄKTA FPLC (GE Healthcare, Amersham Biosciences). SDS-PAGE was performed with 10% gels (Invitrogen) under reducing and non-reducing conditions. For ESI-MS analysis samples were diluted to about 0.1 mg/mL and LC-MS was performed on a Waters Xevo G2XS Qtof instrument (ESI-ToF-MS) coupled to a Waters Acquity UPLC H-Class System using a 2.1 × 50 mm Acquity BEH300 C4 1.7 μm column (Waters). Differential scanning fluorimetry was performed on an Applied Biosystems StepOnePlus RT-PCR instrument. Protein samples were diluted at 2μM in PBS in 40 μL and placed in PCR tubes, assay was performed in triplicates. 5x SYPRO ORANGE (Invitrogen, stock 5000x) was added to samples prior to analysis. For thermal stability measurements, the temperature range spanned from 25°C to 95°C with a scan rate of 1 °C/min. Data analysis was performed in Protein Thermal Shift™ Software version 1.3. The temperature derivative of the melting curve was computed.

### Affinity measurements

Affinity measurements were performed by surface plasmon resonance using BIAcore X100 (BIAcore, GE Healthcare) instrument using a biotinylated CAIX coated streptavidin chip. Samples were injected as serial-dilutions, in a concentration range from 1mM to 62.5nM. Regeneration of the chip was performed by HCl 10 mM.

### *In vitro* biological activities

The biological activity of TNF was determined by incubation with mouse LM fibroblasts, in the presence of 2μg/mL actinomycin D (Sigma-Aldrich). In 96-well plates, cells (20’000 per well) were incubated in medium supplemented with actinomycin D and varying concentrations of recombinant human TNF or IL2-XE114-TNF^mut^. After 24 h at 37C, cell viability was determined with Cell Titer Aqueous One Solution (Promega). Results were expressed as the percentage of cell viability compared to cells treated with actinomycin D only.

The biological activity of IL2 was determined by its ability to stimulate the proliferation of CTLL2 cells. Cells (25’000 per well) were seeded in 96-well plates in the culture medium supplemented with varying concentrations of the fusion proteins. After incubation at 37°C for 48 hours, cell proliferation was determined with Cell Titer Aqueous One Solution (Promega). Results were expressed as the percentage of cell viability compared to untreated cells.

### Flow cytometry

Antigen expression on SKRC52 cells was confirmed by flow cytometry. Cells were centrifuged and washed in cold FACS buffer (0.5% BSA, 2mM EDTA in PBS) and stained with IL2-XE114-TNF^mut^ (final concentration 10μg/mL) and detected with rat anti-IL2 (eBioscience 14-7029-85) followed by staining with anti-rat AlexaFluor488 (Invitrogen A21208). IL2-KSF-TNF^mut^ (specific for an irrelevant antigen) was used as negative control.

### Immunofluorescence studies

Antigen expression was confirmed on ice-cold acetone fixed 8-μm cryostat sections of SKRC52 and CT26-CAIX stained with IL2-XE114-TNF^mut^ and IL2-F8-TNF^mut^ (final concentration 5μg/mL) and detected with rat anti-IL2 (eBioscience 14-7029-85) and anti-rat AlexaFluor488 (Invitrogen A21208). For vascular staining goat anti-CD31 (R&D AF3628) and anti-goat AlexaFluor594 (Invitrogen A11058) antibodies were used. IL2-KSF-TNF^mut^ (specific for an irrelevant antigen) was used as negative control. Slides were mounted with fluorescent mounting medium and analysed with Axioskop2 mot plus microscope (Zeiss).

For ex vivo immunofluorescence analysis, mice were injected with 50-60μg IL2-XE114-TNF^mut^, IL2-F8-TNF^mut^ or IL2-KSF-TNF^mut^ and sacrificed 24 hours after injection. Organs were excised and embedded in cryo-embedding medium (Thermo Scientific) and cryostat section (10μm) were stained using the following antibodies: rat anti-IL2 (eBioscience 14-7029-85) and anti-rat AlexaFluor488 (Invitrogen A21208). For vascular staining goat anti-CD31 (R&D AF3628) and anti-goat AlexaFluor594 (Invitrogen A11058) antibodies were used. Slides were mounted with fluorescent mounting medium and analysed with Axioskop2 mot plus microscope (Zeiss).

### Mice therapy studies

Mice were monitored daily and tumor volume was measured with a caliper (volume = length × width^2^ × 0.5). When tumors reached a suitable volume (approx. 70–100 mm^3^), mice were injected into the lateral tail vein with the pharmacological agents. Fusion proteins were dissolved in PBS, also used as negative control, and administered at 30μg four times every 24 hours. Results are expressed as tumor volume in mm^3^ ± SEM and % mean body weight change ± SEM. For therapy experiments n = 5 mice/group.

### Non-human primate study

The non-human primate study was performed in accordance with the Directive 2010/63/UE of the European parliament and of the council of 22 September 2010 for the protection of animals used for scientific purposes. Approval for the test site of experimentation: No. E 18-023-01. A total of 3 male naïve Macaca fascicularis (Cynomolgus monkey, Old Java monkey), approximately 30 months at the time of allocation and estimated to weigh between 2.5 to 4.0 kg were used in this study. Test items were administered by bolus intravenous injection in the cephalic or saphenous vein, at a dose volume of 0.5 mL/kg body weight (corresponding to 0.1mg/kg), over a period of approximately 30 seconds. A flush with 1 mL of physiological saline was administered at the end of the bolus injection. The dose was administered to each animal on the basis of the body weight measured on the day of administration. Blood samples of approximately 1 mL each were collected from the saphenous or cephalic vein (alternatively from other blood vessels) of all animals at approximately the following 7 time points: before dosing and at 1, 15 and 30 minutes and 1, 2, and 4 hours after treatment. Samples were transferred into serum separator tubes, kept for 30 minutes in an upright position then centrifuged at room temperature (2500 g for 10 minutes) and the serum divided into 2 polypropylene tubes. Tubes were frozen within 90 minutes post blood sampling and stored at −80±10°C.

### Pharmacokinetics analysis

Fusion protein concentrations in serum were assessed by AlphaLISA. Briefly, Streptavidin Donor Beads were coated with biotinylated antigen (CAIX for IL2-XE114-TNF^mut^ or EDA for IL2-F8-TNF^mut^). Acceptor Beads coated with an anti-TNF antibody were used for detection.

### Statistical analysis

Data were analysed using Prism 7.0 (GraphPad Software, Inc.). Differences in tumor volume between therapeutic groups (until day 14, when n = 5) were evaluated with the two-way ANOVA followed by Bonferroni as post-test. P < 0.05 was considered statistically significant. (*: P < 0.05, **: P < 0.01, ***: P < 0.001, ****: P < 0.0001).

## Results

**Figure 1A** depicts a schematic representation of a fully-human fusion protein (termed IL2-XE114-TNF^mut^), featuring a sequential arrangement of IL2, a scFv fragment specific to CAIX (named XE114 (31)) and TNF [**Supplementary Figure 1**]. The protein arrangement is reminiscent of the one previously described for the murine fusion protein IL2-F8-TNF^mut^ (6,27) [**Supplementary Figure 2**], but here we used human payloads in order to facilitate clinical translational activities. The TNF moiety was de-potentiated by a single amino acid substitution (R108A), in order to achieve a similar cytokine activity for both IL2 and TNF moieties. ScFv(XE114) has previously been shown to recognize human CAIX with high affinity and kinetic stability (31,35). IL2-XE114-TNF^mut^ was expressed in mammalian cells and could be purified on Protein A, since the scFv moiety featured a VH domain of the VH3 family (36,37). The product formed stable non-covalent homotrimers in solution [**Figure 1B**], as TNF is a trimeric protein, and bound avidly to the cognate antigen in BIAcore assays [**Figure 1C**]. The protein was mainly in a non-glycosylated form, but approx. 10% of IL2-XE114-TNF^mut^ exhibited a molecular weight increase of 657 Dalton, as a result of O-linked glycosylation [**Figure 1D**]. The product retained intact IL2 activity in an *in vitro* lymphocyte proliferation assay [**Figure 1E**], while TNF potency was reduced by approximately 10-fold, as a result of a single amino acid substitution (6) [**Figure 1F**]. A single band could be detected in SDS-PAGE analysis, both in reducing and in non-reducing conditions [**Figure 1G**]. IL2-XE114-TNF^mut^ bound to SKRC52 renal cell carcinoma cell lines more intensely than IL2-KSF-TNF^mut^ [**Figure 1H**], which was specific to hen egg lysozyme and was chosen as a negative control of irrelevant specificity in the mouse (38) [**Supplementary Figure 3**]. A multi-step denaturation profile was observed by differential scanning fluorimetry, with a first transition at 49.5°C [**Figure 1J**].

**Figure 1:**
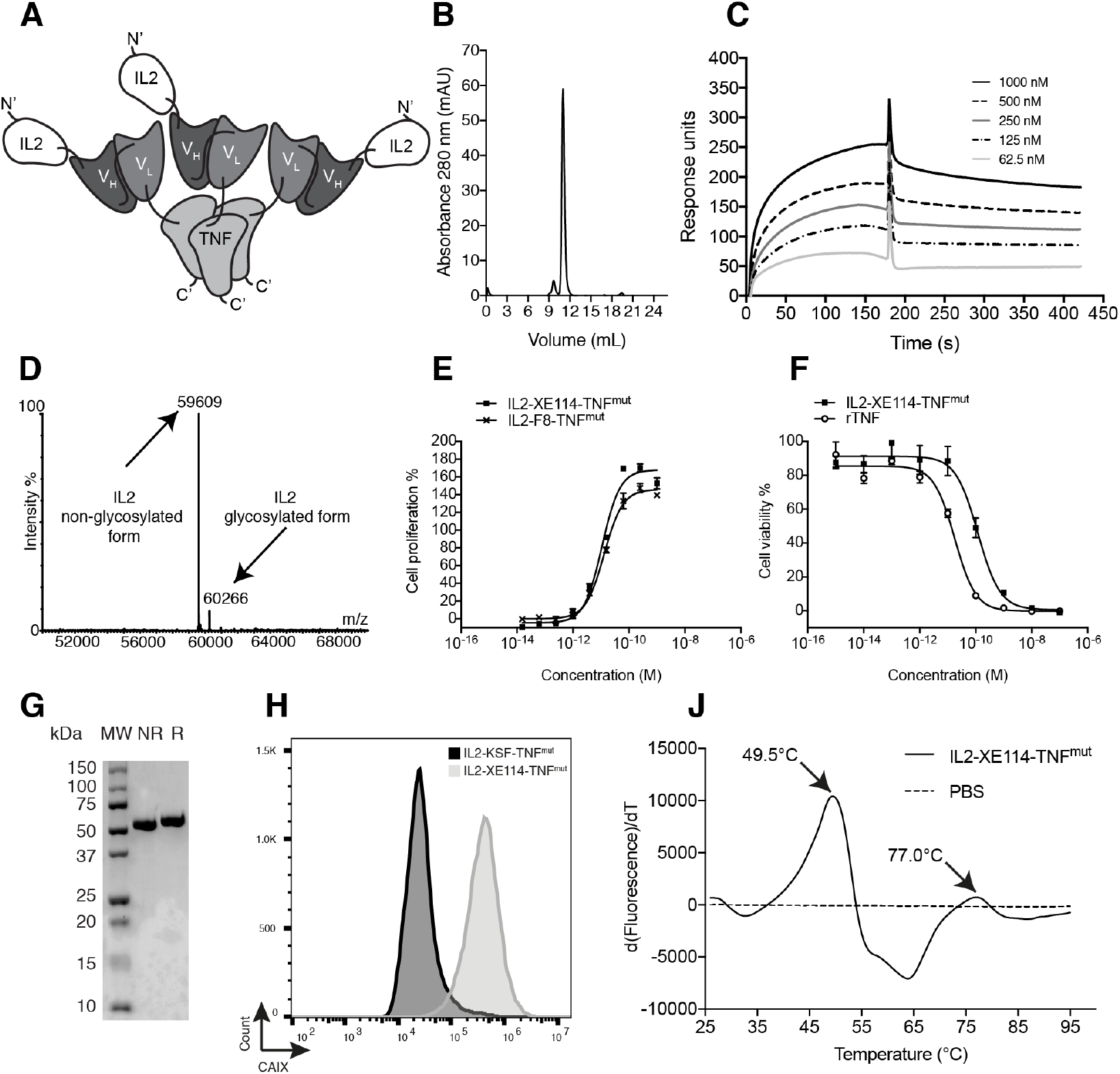
Biochemical characterization of IL2-XE114-TNF^mut^. [**A**] Schematic representation of the domain assembly of IL2-XE114-TNF^mut^ in noncovalent homotrimeric format. [**B**] Size exclusion chromatography profile. [**C**] BIAcore analysis on CAIX-coated sensor chip. [**D**] ESI-MS profile. [**E**] IL2 bioactivity assay on CTLL2 cells. [**F**] TNF bioactivity assay on L-M fibroblasts. [**G**] SDS-PAGE analysis; MW: molecular weight, NR: non-reducing conditions, R: reducing conditions. [**H**] Flow cytometric evaluation of CAIX expression by SKRC52 cells, detected with IL2-XE114-TNF^mut^. [**J**] Differential Scanning Fluorimetry on IL2-XE114-TNF^mut^.

The tumor-homing properties of IL2-XE114-TNF^mut^ were first studied in nude mice, bearing subcutaneously-grafted human SKRC52 renal cell carcinomas [**Figure 2**]. Organs were examined 24 hours after intravenous administration of 60 μg of fusion protein, using an immunofluorescence procedure for the detection of the IL2 moiety. A homogenous antigen expression pattern was observed in an *in vitro* analysis of tumor sections. However, *ex vivo*, IL2-XE114-TNF^mut^ mainly localized to perivascular tumor cells and failed to homogenously stain the CAIX-positive tumor mass. Similar targeting behaviors have previously been reported for other antibody products, directed against cell surface tumor antigens (31,39). No detectable antibody uptake could be seen in relevant normal organs [**Figure 2**]. By contrast, no tumor staining and no tumor uptake could be observed for the IL2-KSF-TNF^mut^ negative control protein [**Figure 2B**].

**Figure 2:**
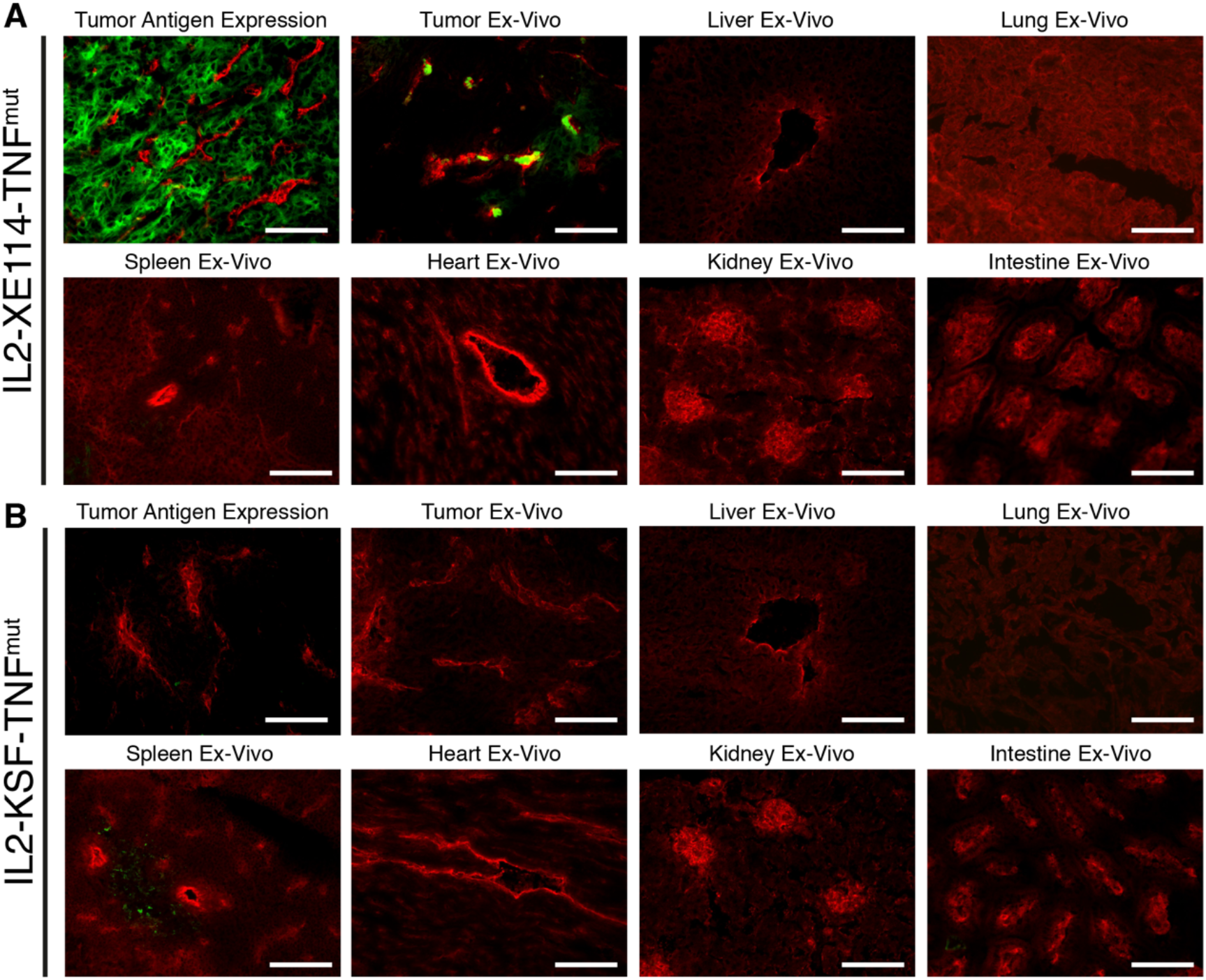
Antigen expression and tumor targeting properties of IL2-XE114-TNF^mut^. Microscopic fluorescence analysis of CAIX expression on SKRC52 tumor sections [**A, B upper left**] detected with IL2-XE114-TNF^mut^ or IL2-KSF-TNF^mut^ (negative control). Microscopic fluorescence analysis of organs from SKRC52 tumor bearing mice, 24 hours after intravenous administration of IL2-XE114-TNF^mut^ [**A**] or IL2-KSF-TNF^mut^ [**B**]. Cryosections were stained with anti-IL2 (green, AlexaFluor 488) and anti-CD31 (red, AlexaFluor 594). 20x magnification, scale bars = 100μm.

In order to get a finer characterization of the tumor-targeting properties of our products, we compared IL2-XE114-TNF^mut^ and IL2-F8-TNF^mut^ (an analogue specific to the alternatively-spliced EDA domain of fibronectin) [**Supplementary Figure 4**] in mice bearing murine CT26 tumors, that had been stably-transfected for CAIX expression on the cell surface (30). Both products exhibited a preferential accumulation at the tumor site, while IL2-KSF-TNF^mut^ failed to localize to the neoplastic mass. However, the IL2-F8-TNF^mut^ yielded a more homogenous pattern of tumor uptake compared to IL2-XE114-TNF^mut^, in spite of the fact that CAIX was strongly expressed on all tumor cells [**Figure 3**].

**Figure 3:**
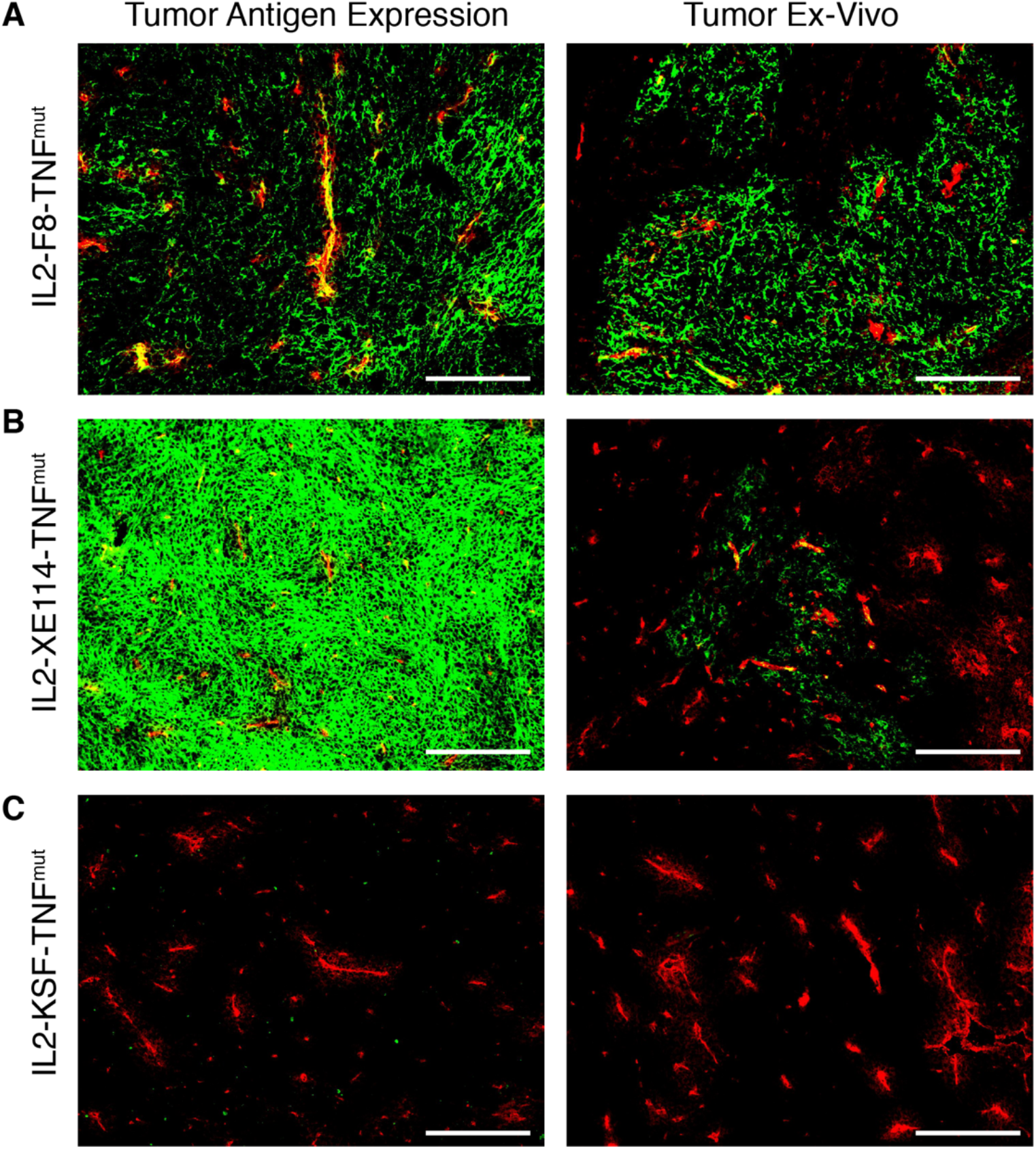
Antigen expression and tumor targeting properties of IL2-XE114-TNF^mut^. Microscopic fluorescence analysis of CAIX expression on CT26-CAIX tumor sections [**A, B, C left**] detected with IL2-F8-TNF^mut^, IL2-XE114-TNF^mut^ or IL2-KSF-TNF^mut^ (negative control). Microscopic fluorescence analysis of tumors from CT26-CAIX tumor bearing mice, 24 hours after intravenous administration of IL2-F8-TNF^mut^ [**A**], IL2-XE114-TNF^mut^ [**B**] or IL2-KSF-TNF^mut^ [**C**]. Cryosections were stained with anti-IL2 (green, AlexaFluor 488) and anti-CD31 (red, AlexaFluor 594). 10x magnification, scale bars = 200μm.

We tested the therapeutic activity of IL2-XE114-TNF^mut^ and IL2-F8-TNF^mut^ in nude mice bearing CAIX-transfected CT26 tumors [**Figure 4**]. Since the human TNF moiety is only partly active in mice, we also studied the therapeutic activity of the mIL2-XE114-mTNF^mut^ and mIL2-F8-mTNF^mut^ analogues, bearing an attenuated version of murine TNF (6,27) [**Supplementary Figure 2, 5**]. A tumor-growth retardation was observed in this model, which lacked a functional set of T lymphocytes, but still retained natural killer (NK) cells [**Figure 4**]. All products caused a transient reduction in body weight at the dose used (30 μg), which was below the 10% threshold.

**Figure 4:**
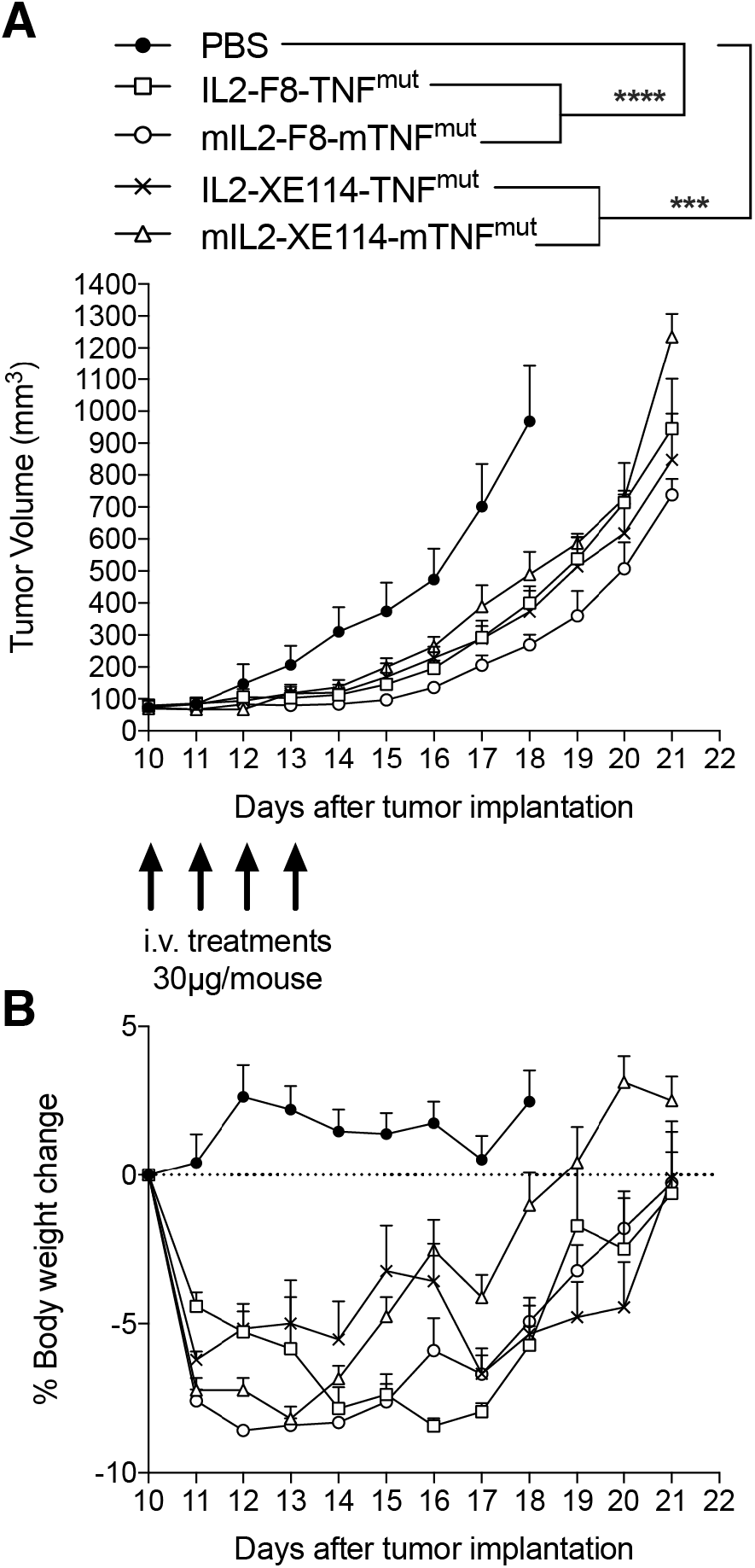
Therapeutic performance of IL2-XE114-TNF^mut^ in BALB/c nude mice bearing CT26-CAIX tumors in comparison with IL2-F8-TNF^mut^ and their murine surrogates (mIL2-XE114-mTNF^mut^, mIL2-F8-mTNF^mut^). Treatment started when tumors reached a volume of 70–100 mm^3^, mice were injected intravenously four times every 24 hours with 30μg of fusion proteins. Results are expressed as tumor volume in mm^3^ ± SEM [**A**] and % mean body weight change ± SEM [**B**]. For therapy experiments n = 5 mice/group. n = 4 for group IL2-XE114-TNF^mut^ from day 15. n = 4 for groups mIL2-XE114-mTNF^mut^ and mIL2-F8-mTNF^mut^ from day 19. n = 3 for group mIL2-F8-mTNF^mut^ from day 20.

Finally, we compared the pharmacokinetic profiles of IL2-XE114-TNF^mut^ and IL2-F8-TNF^mut^ in *Cynomolgus* monkey, after a single intravenous administration (0.1mg/kg) [**Figure 5**]. IL2-F8-TNF^mut^ showed a biphasic clearance profile, with a rapid loss of ~ 2/3 of the protein from circulation, followed by a slower elimination phase. By contrast, IL2-XE114-TNF^mut^ exhibited a slower clearance profile, with a half-life of approximately 2 hours. This pharmacokinetic profile is similar to the one that we have previously observed for other antibody-cytokine fusion proteins, which have progressed to advanced clinical trials (13,18,40,41).

**Figure 5:**
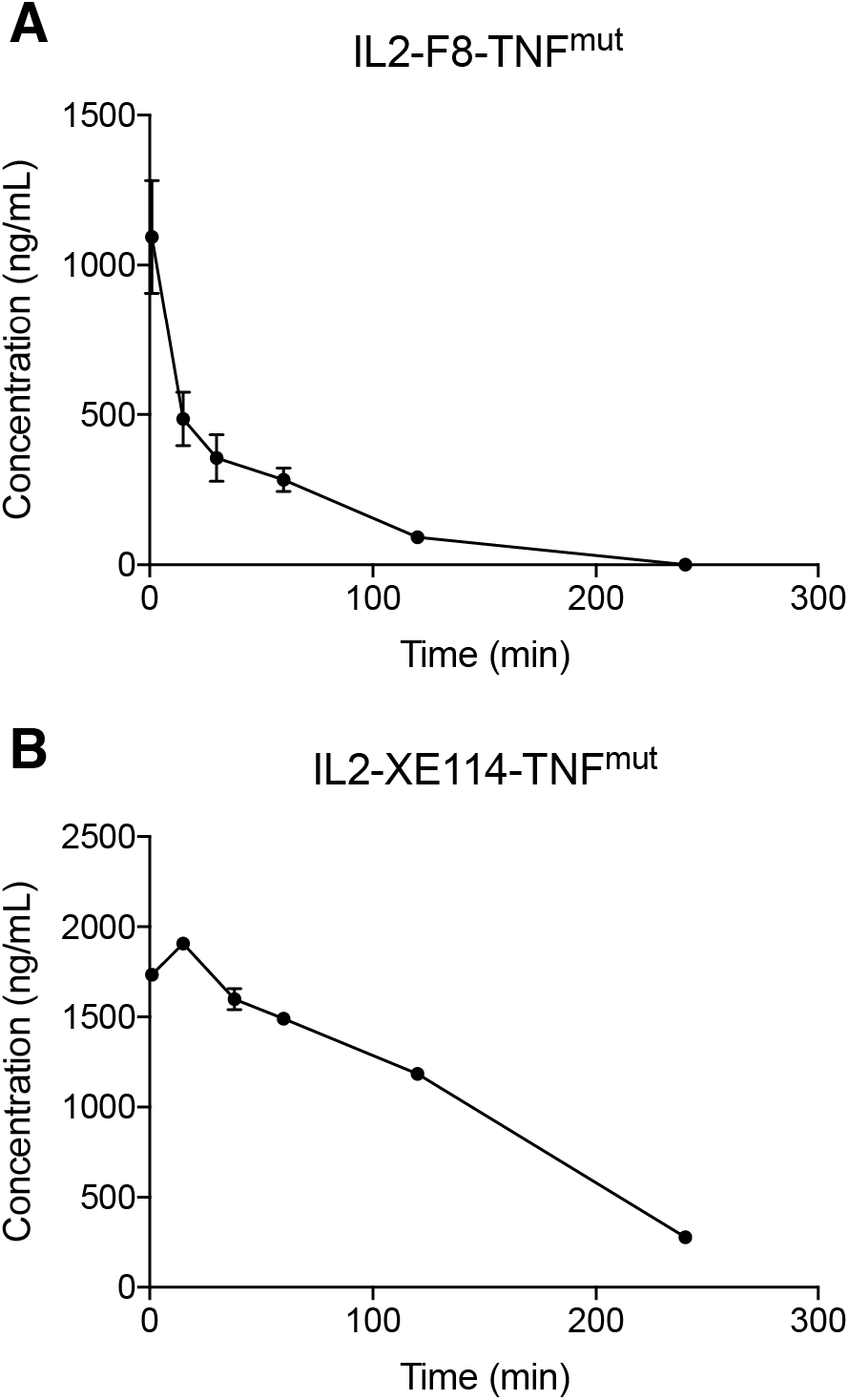
Pharmacokinetics in cynomolgus monkeys. Pharmacokinetics were evaluated in one cynomolgus monkey per group injected once at the dose of 0.1mg/kg of IL2-F8-TNF^mut^ [**A**] or IL2-XE114-TNF^mut^ [**B**]. Blood samples were collected before dosing and at 1, 15 and 30 minutes and 1, 2, and 4 hours after treatment.

## Discussion

In this work we have described the generation, the *in vitro* characterization and the *in vivo* validation of a novel “potency-matched dual-cytokine antibody fusion protein” based on an anti-CAIX antibody fragment simultaneously fused to human IL2 and to a mutant version of human TNF. The product, named IL2-XE114-TNF^mut^, could be expressed in mammalian cells and purified to homogeneity. The fusion protein was able to recognize CAIX both *in vitro* and *in vivo*, when tested on a human renal cell carcinoma cell line (SKRC52, which naturally express the antigen) and in a murine colon carcinoma cell line (CT26-CAIX, which had been transfected with the human antigen).

We have previously reported that the intralesional administration of two immunocytokine products (L19-IL2 and L19-TNF) was able to induce cancer remission both in mouse models of cancer (24) and in patients with stage IIIB/C melanoma (22). However, from an industrial prospective, the development of combination products may cause a duplication of activities and costs. By contrast, the opportunity of incorporate two cytokine payloads into the same antibody moiety may facilitate industrial development.

IL2- and TNF-based products have shown to be synergistically active against various type of malignancies (4,6,8,22–25,27,42) by two distinct complementary mechanism of actions. On one hand, TNF is capable of inducing hemorrhagic necrosis and apoptosis of the tumor endothelial cells and also of cancer cells (4,6,25,43,44). On the other end, IL2 is able to promote a selective boosting of T cell and NK cell activity against cancer cells (6,24,26,45).

The therapeutic activity of IL2-XE114-TNF^mut^ was evaluated in immunocompromised mice bearing human CAIX-transfected CT26 murine tumors. This mouse model, which lacked a functional set of T lymphocytes, but still retained NK cells, was chosen in order to avoid an immune response against the transfected human antigen. This may explain the reason why only an initial tumor growth inhibition was observed. We had previously reported that depletion studies in immunocompetent mice treated with IL2-F8-TNF^mut^ revealed a prominent role of CD4+ and CD8+ T cells in the cancer remission process (6).

Antibodies against cell surface antigens can recognize their cognate antigen with exquisite specificity, but often their penetration into solid malignancies can be suboptimal as a result of a slow extravasation rate due to their relatively large size (39,46). Indeed, IL2-XE114-TNF^mut^ was found to preferentially localize to perivascular cells *in vivo*. Similar findings had been reported for antibodies specific to HER2 (39). Interestingly, it has been shown that a small molecule against CAIX could penetrate CAIX-positive tumors more efficiently compared to an antibody against the same target (31).

In spite of a suboptimal penetration in neoplastic lesions, IL2-XE114-TNF^mut^ revealed an acceptable clearance profile, with a half-life of approximately 2 hours in monkeys, which was more favorable compared to the one of IL2-F8-TNF^mut^. This pharmacokinetic profile is comparable to the one previously observed for other cytokine-fusion proteins based on antibody-fragments (40,47). By contrast, IL2-F8-TNF^mut^ exhibited a rapid clearance from circulation already at early time-points, possibly indicating a trapping of the F8 antibody in the liver, which was already observed for other F8-based immunocytokines (48).

Renal cell carcinoma (RCC) represents a rare condition that account for approx. 2% of cancer deaths worldwide (49). The most prominent subtype of RCC (about 70%) is clear cell (ccRCC) (50). In most cases (70%) the tumor is confined to the kidney, but it may disseminate to regional lymph nodes and to visceral organs (50). Surgical resection is the primary treatment option for stage I-III RCC, but postsurgical recurrence is observed with a 5-year relapse rate of 30-40% in patients with stage II or III RCC (51). In the event of recurrence patients are typically treated with conventional chemotherapy (e.g. sunitinib), high dose IL2 or immune-checkpoint inhibitors (e.g. nivolumab and ipilimumab) (52,53). A phase III clinical trials in patients with advanced renal-cell carcinoma showed that the combination of nivolumab and ipilimumab could increase the overall survival compared to sunitinib alone (53). However, this efficacy was observed only in a small portion of patients, for this reason novel therapeutics for the treatment of RCC may be required.

CAIX is strongly expressed in the majority of ccRCC. The antigen has been extensively validated as a target for ccRCC in preclinical studies and several antibodies against this antigen have been developed by our group and others (54,55). An humanized monoclonal antibody, named G250 (56) has been used to validate CAIX as a cancer target by nuclear medicine in clinical studies (50,57). The XE114 antibody fragment used in this study is a fully-human high-affinity antibody (31), which may be considered for clinical development of CAIX-targeted based therapeutics.

The product presented in this study (IL2-XE114-TNF^mut^) was able to target CAIX in tumor bearing mice and showed a therapeutic effect in immunocompromised animals. Moreover, the favorable pharmacokinetic profile in monkey provide a rational for future clinical investigation. The targeted delivery of cytokine payloads to cell surface antigens may represent a valid alternative to the anchoring of pro-inflammatory payloads to the tumor extracellular matrix. To a certain extent, an immunocytokine able to selectively localize to tumor cell membrane may represent a functional equivalent to a “bispecific antibody” and may be capable of cross-link a tumor cell with a leukocyte (e.g. NK cell, T cell), which displays the cognate cytokine receptor on its surface.

## Acknowledgments

We would like to thank Dr. Samuele Cazzamalli for helpful discussions and we gratefully acknowledge Lisa Nadal and Riccardo Corbellari for their help with experimental procedures.

## Funding

We gratefully acknowledge funding from ETH Zürich and the Swiss National Science Foundation (Grant Nr. 310030_182003/1). This project has received funding from the European Research Council (ERC) under the European Union’s Horizon 2020 research and innovation program (grant agreement 670603).

## Conflict of interest

Dario Neri is co-founder, shareholder and member of the board of Philogen, a company working on antibody therapeutics. The authors declare no additional conflict of interest.

## Authors contributions

RD, DN: conception and design; development of methodology; acquisition, analysis and interpretation of data; writing, review and revision of the manuscript. AV, SG: analysis and interpretation of data. BG, TO, BZ, AS, GB, MM: technical support. RD, DN: study supervision.

