## Supplementary Information for "A novel fully-human potency-matched dual cytokine-antibody fusion protein targets carbonic anhydrase IX in renal cell carcinomas"

**Supplementary Material**

IL2 – 12aa linker – $V_H$ – 14aa linker – $V_L$ – 15aa linker – $TNF^{mut}$

APTSSSTKKTQLQLEHLLLDLQMILNGINNYKNPKLTRMLTFKFYMPKKATELKHLQCLEEE
LKPLEEVLNLAQSKNFHLRPRDLISNINVIVLELKGSETTFMCEYADETATIVEFLNRWITF
CQSIISTLTGDGSSGGSGGASEVQLLESGGGLVQPGGSLRLSCAASGFTFSSYAMSWVRQAP
GKGLEWVSAIDGSGGSTYYADSVKGRFTISRDNSKNTLYLQMNSLRAEDTAVYYCVKGPPVF
DYWGQGTLVTVSSGGGGSGGGGSGGGGSSELTQDPAVSVALGQTVRITCQGDSLRSYYASWY
QQKPGQAPVLVIYGKNNRPSGIPDRFSGSSSGNTASLTITGAQAEDEADYYCQSSKWSWDPV
VFGGGTKLTVLGSSSSGSSSSGSSSSGVRSSSRTPSDKPVAHVVANPQAEGQLQWLNRAANA
LLANGVELRDNQLVVPSEGLYLIYSQVLFKGQGCPSTHVLLTHTISRIAVSYQTKVNLLSAI
KSPCQRETPEGAEAKPWYEPIYLGGVFQLEKGDRLSAEINRPDYLDFAESGQVYFGIIAL

**Supplementary Figure 1:** Amino acid sequence of IL2-XE114-$TNF^{mut}$. Starting from the N-terminus: human IL2, the XE114 antibody in scFv format and human TNF bearing the R431A mutation, corresponding to the position 32 in the soluble form of TNF.

IL2 – 12aa linker – V$_H$ – 14aa linker – V$_L$ – 15aa linker – TNF$^{mut}$

```
APTSSSTSSSTAEAQQQQQQQQQQQQHLEQLLMDLQELLSRMENYRNLKLPRMLTFKFYLPK
QATELKDLQCLEDELGPLRHVLDLTQSKSFQLEDAENFISNIRVTVVKLKGSDNTFECQFDD
ESATVVDFLRRWIAFCQSIISTSPQGDGSSGGSGGASEVQLLESGGGLVQPGGSLRLSCAAS
GFTFSLFTMSWVRQAPGKGLEWVSAISGSGGSTYYADSVKGRFTISRDNSKNTLYLQMNSLR
AEDTAVYYCAKSTHLYLFDYWGQGTLVTVSSGGGGSGGGGSGGGGEIVLTQSPGTLSLSPGE
RATLSCRASQSVSMPFLAWYQQKPGQAPRLLIYGASSRATGIPDRFSGSGSGTDFTLTISRL
EPEDFAVYYCQQMRGRPPTFGQGTKVEIKSSSSGSSSSGSSSSGLRSSSQNSSDKPVAHVVA
NHQVEEQLEWLSQWANALLANGMDLKDNQLVVPADGLYLVYSQVLFKGQGCPDYVLLTHTVS
RFAISYQEKVNLLSAVKSPCPKDTPEGAELKPWYEPIYLGGVFQLEKGDQLSAEVNLPKYLD
FAESGQVYFGVIAL
```

**Supplementary Figure 2:** Amino acid sequence of mIL2-F8-mTNF$^{mut}$. Starting from the N-terminus: murine IL2, the F8 antibody in scFv format and murine TNF bearing the R448W mutation, corresponding to the position 32 in the soluble form of TNF.

IL2 – 12aa linker – $V_H$ – 14aa linker – $V_L$ – 15aa linker – $TNF^{mut}$

APTSSSTKKTQLQLEHLLLDLQMILNGINNYKNPKLTRMLTFKFYMPKKATELKHLQCLEEE
LKPLEEVLNLAQSKNFHLRPRDLISNINVIVLELKGSETTFMCEYADETATIVEFLNRWITF
CQSIISTLTGDGSSGGSGGASEVQLLESGGGLVQPGGSLRLSCAASGFTFSSYAMSWVRQAP
GKGLEWVSAISGSGGSTYYADSVKGRFTISRDNSKNTLYLQMNSLRAEDTAVYYCAKSPKVS
LFDYWGQGTLVTVSSGGGGSGGGGSGGGGSSELTQDPAVSVALGQTVRITCQGDSLRSYYAS
WYQQKPGQAPVLVIYGKNNRPSGIPDRFSGSSSGNTASLTITGAQAEDEADYYCNSSPLNRL
AVVFGGGTKLTVLGSSSSGSSSSGSSSSGVRSSSRTPSDKPVAHVVANPQAEGQLQWLNRAA
NALLANGVELRDNQLVVPSEGLYLIYSQVLFKGQGCPSTHVLLTHTISRIAVSYQTKVNLLS
AIKSPCQRETPEGAEAKPWYEPIYLGGVFQLEKGDRLSAEINRPDYLDFAESGQVYFGIIAL

**Supplementary Figure 3:** Amino acid sequence of IL2-KSF-$TNF^{mut}$. Starting from the N-terminus: human IL2, the KSF antibody in scFv format and human TNF bearing the R433A mutation, corresponding to the position 32 in the soluble form of TNF.

IL2 – 12aa linker – $V_H$ – 14aa linker – $V_L$ – 15aa linker – $TNF^{mut}$

APTSSSTKKTQLQLEHLLLDLQMILNGINNYKNPKLTRMLTFKFYMPKKATELKHLQCLEEE
LKPLEEVLNLAQSKNFHLRPRDLISNINVIVLELKGSETTFMCEYADETATIVEFLNRWITF
CQSIISTLTGDGSSGGSGGASEVQLLESGGGLVQPGGSLRLSCAASGFTFSLFTMSWVRQAP
GKGLEWVSAISGSGGSTYYADSVKGRFTISRDNSKNTLYLQMNSLRAEDTAVYYCAKSTHLY
LFDYWGQGTLVTVSSGGGGSGGGGSGGGGEIVLTQSPGTLSLSPGERATLSCRASQSVSMPF
LAWYQQKPGQAPRLLIYGASSRATGIPDRFSGSGSGTDFTLTISRLEPEDFAVYYCQQMRGR
PPTFGQGTKVEIKSSSSGSSSSGSSSSGVRSSSRTPSDKPVAHVVANPQAEGQLQWLNRAAN
ALLANGVELRDNQLVVPSEGLYLIYSQVLFKGQGCPSTHVLLTHTISRIAVSYQTKVNLLSA
IKSPCQRETPEGAEAKPWYEPIYLGGVFQLEKGDRLSAEINRPDYLDFAESGQVYFGIIAL

**Supplementary Figure 4:** Amino acid sequence of IL2-F8-$TNF^{mut}$. Starting from the N-terminus: human IL2, the F8 antibody in scFv format and human TNF bearing the R432A mutation, corresponding to the position 32 in the soluble form of TNF.

IL2 – 12aa linker – $V_H$ – 14aa linker – $V_L$ – 15aa linker – $TNF^{mut}$

APTSSSTSSSTAEAQQQQQQQQQQQQHLEQLLMDLQELLSRMENYRNLKLPRMLTFKFYLPK
QATELKDLQCLEDELGPLRHVLDLTQSKSFQLEDAENFISNIRVTVVKLKGSDNTFECQFDD
ESATVVDFLRRWIAFCQSIISTSPQGDGSSGGSGGASEVQLLESGGGLVQPGGSLRLSCAAS
GFTFSSYAMSWVRQAPGKGLEWVSAIDGSGGSTYYADSVKGRFTISRDNSKNTLYLQMNSLR
AEDTAVYYCVKGPPVFDYWGQGTLVTVSSGGGGSGGGGSGGGGSSELTQDPAVSVALGQTVR
ITCQGDSLRSYYASWYQQKPGQAPVLVIYGKNNRPSGIPDRFSGSSSGNTASLTITGAQAED
EADYYCQSSKWSWDPVVFGGGTKLTVLGSSSSGSSSSGSSSSGLRSSSQNSSDKPVAHVVAN
HQVEEQLEWLSQWANALLANGMDLKDNQLVVPADGLYLVYSQVLFKGQGCPDYVLLTHTVSR
FAISYQEKVNLLSAVKSPCPKDTPEGAELKPWYEPIYLGGVFQLEKGDQLSAEVNLPKYLDF
AESGQVYFGVIAL

**Supplementary Figure 5:** Amino acid sequence of mIL2-XE114-$mTNF^{mut}$. Starting from the N- terminus: murine IL2, the XE114 antibody in scFv format and murine TNF bearing the R447W mutation, corresponding to the position 32 in the soluble form of TNF.
